# Building computational benchmarks: an Omnibenchmark reimplementation of a single-cell preprocessing pipeline evaluation

**DOI:** 10.64898/2026.05.01.722166

**Authors:** Atreya Choudhury, Tom Kitak, Ben Carrillo, Pauline Busch, Martin Emons, Samuel Gunz, Maruša Koderman, Siyuan Luo, Izaskun Mallona, Aidan Meara, David Wissel, Mark D. Robinson

## Abstract

**Background:** In the past few years, we have seen a veritable surge in single-cell (e.g., RNA-sequencing) techniques and datasets, enabling increasingly detailed characterization of cellular heterogeneity across tissues and conditions. This surge in single-cell techniques has been complemented by a large number of analysis frameworks and pipelines, and a large parameter space and researcher degrees of freedom to use them. Many neutral benchmarks have been presented for various computational tasks, but most make design decisions that render them incompatible with each other, e.g., through the use of different datasets, metrics, or parameter sets.

**Results:** We showcase Omnibenchmark, a recently developed framework to build reproducible, extensible, and standardized method comparisons for an arms-length benchmark that evaluates five single-cell RNA sequencing pipelines, spanning filtering, normalization, dimensionality reduction, and clustering, on three datasets. The framework facilitates the broad investigation of pipelines used in single-cell data analysis while highlighting how the process of building benchmarks can be streamlined and unified. This standardization enables benchmarks to be extended in several directions, including broader parameter sweeps, comparisons across software versions and architectures, isolation of pipeline steps, and integration of additional pipelines, datasets, and metrics. As an example, our harmonization of pipelines highlighted that tools implement the (clustering) resolution parameter differently.

**Conclusions:** Omnibenchmark provides a reproducible, extensible, and standardized approach for bench-marking. By unifying the process of benchmark construction, it can facilitate more consistent method comparisons and provides a foundation for broader, more flexible, and more sustainable benchmarking efforts.

## Background

In the past decade, there has been an explosion in both the scale and diversity of single-cell datasets. While single-cell RNA sequencing (scRNA-seq) remains the most widely-adopted, it now sits within a broader ecosystem of adjacent single-cell technologies, encompassing protein-centric assays such as CyTOF [1], multi-modal measurements that couple RNA with surface proteins (CITE-seq [2]), chromatin accessibility profiling via scATAC-seq [3], CRISPR-based perturbation screens coupled with RNA-seq [4], and other emerging single-cell multiomics methods that jointly profile molecular layers within the same cell (or across matched cells) [5, 6]. In parallel with the growth in datasets, there has been a rapid proliferation of computational methods for single-cell analysis. The scrna-tools database has documented an accelerating expansion of the method landscape [7, 8], revealing more than 1500 tools, with visualization, dimensionality reduction and clustering being the most-studied tasks. While scientifically valuable, the abundance of choices makes it challenging for practitioners to assemble pipelines with predictable behavior.

The evaluation of the components of a typical pipeline in isolation is a non-trivial task. Upstream choices may modify the measured impact of the component under consideration. The quality and stability of such analyses reflect the cumulative choices made throughout the multiple steps in a data analysis pipeline. Meanwhile, the community has invested heavily in modular and interoperable scRNA-seq analysis frameworks (`scanpy` [9], `osca` [10], `scrapper` [11], `seurat` [12]), alongside broader attempts to standardize data representations (`anndata` [13], `SingleCellExperiment` [10]), workflows (`snakemake` [14], `nextflow` [15]), and benchmarking infrastructures (Open Problems [16], Omnibenchmark [17], OpenEBench [18]). At the same time, increasing dataset sizes and growing interest in interactive analysis have motivated the adoption of scalable computing approaches, including GPU-accelerated libraries (`rapids-singlecell` [19]) or via efficient compression techniques (`BPCells` [20]). In particular, GPU acceleration can dramatically reduce runtimes for computational bottlenecks such as nearest-neighbor graph construction, dimensionality reduction, and clustering, enabling analyses that would otherwise be impractical at atlas scale. On the other hand, bit packing strategies allow analysis of large datasets (e.g., *>*10M cells) on laptop-level hardware. Nonetheless, these performance gains also introduce new sources of variability: upstream computational choices—such as the specific PCA implementation, the handling of sparse matrices, or the floating-point precision used on GPUs influence the representation of the data and can alter the results [21]. Additionally, some implementations have specific hardware requirements.

After a decade of analyzing scRNA-seq data, the field now contains many neutral benchmarks, where evaluation is decoupled from tool development and aims to provide guidance that generalizes beyond a single method family. There are also best-practice efforts that synthesize lessons learned and propose recommended workflows [22, 23], while acknowledging that best practices may evolve as technologies change and as new empirical evidence accumulates. Adjacent to this, recent meta-analyses highlight persistent shortcomings in how benchmarks are conducted. A review of *>*60 single-cell benchmarks found that, although many researchers make code and data available, they are often difficult to extend; missing artifacts, incomplete provenance, unavailable simulation code, and fragile software environments were recurring barriers to extensibility [24]. Similarly, a systematic assessment of *>*280 scRNA-seq methodological studies (130 benchmarks) argued that the rapid growth of benchmarking has not been accompanied by comparable progress in standardization, reporting, and sustainability, limiting the long-term value of many efforts [25].

In this work, we built a standardized benchmark, `scrna-bench`, via Omnibenchmark [17] to focus partially on pipeline performance (both speed and accuracy) for scRNA-seq data analysis, while keeping the scope broad enough to still allow a deep and actionable treatment of critical parts (e.g., parameterization) of the workflow. Here, we leverage a recent “end-to-end pipeline” benchmark as an initial canvas [21]. While the results are interesting on their own, we also view this benchmark as a draft template that can be seeded by interested researchers and iteratively improved through a broader community effort. At the same time, it offers not only standardization and the ability to scrutinize important details, but also the opportunity to isolate components for further study. More broadly, the strategy we pursue here is not specific to scRNA-seq: analogous standardized and extensible benchmarking templates could benefit many areas of computational biology (and other computational subfields) by enabling open source modules that can be reused, reanalyzed, and continuously updated as methods and datasets evolve.

## Results and discussion

The original study [21] compared scRNA-seq pipelines for scalability and accuracy on three datasets and emphasized that algorithmic and infrastructural choices drive performance. Starting from the original authors’ code, we added further structure via a benchmark plan (see topology in Figure 1A), injected code snippets that harmonized inputs and outputs for all stages of the benchmark (datasets, pipelines, metrics), and exposed commonly-tuned parameters to explore a broader grid of the parameter space. We note also that the 5 pipelines share considerable methodology (e.g., normalization, feature selection, PCA, UMAP, Louvain/Leiden graph-based clustering) but each differ in philosophical choices as well as implementation details.

**Fig. 1.**
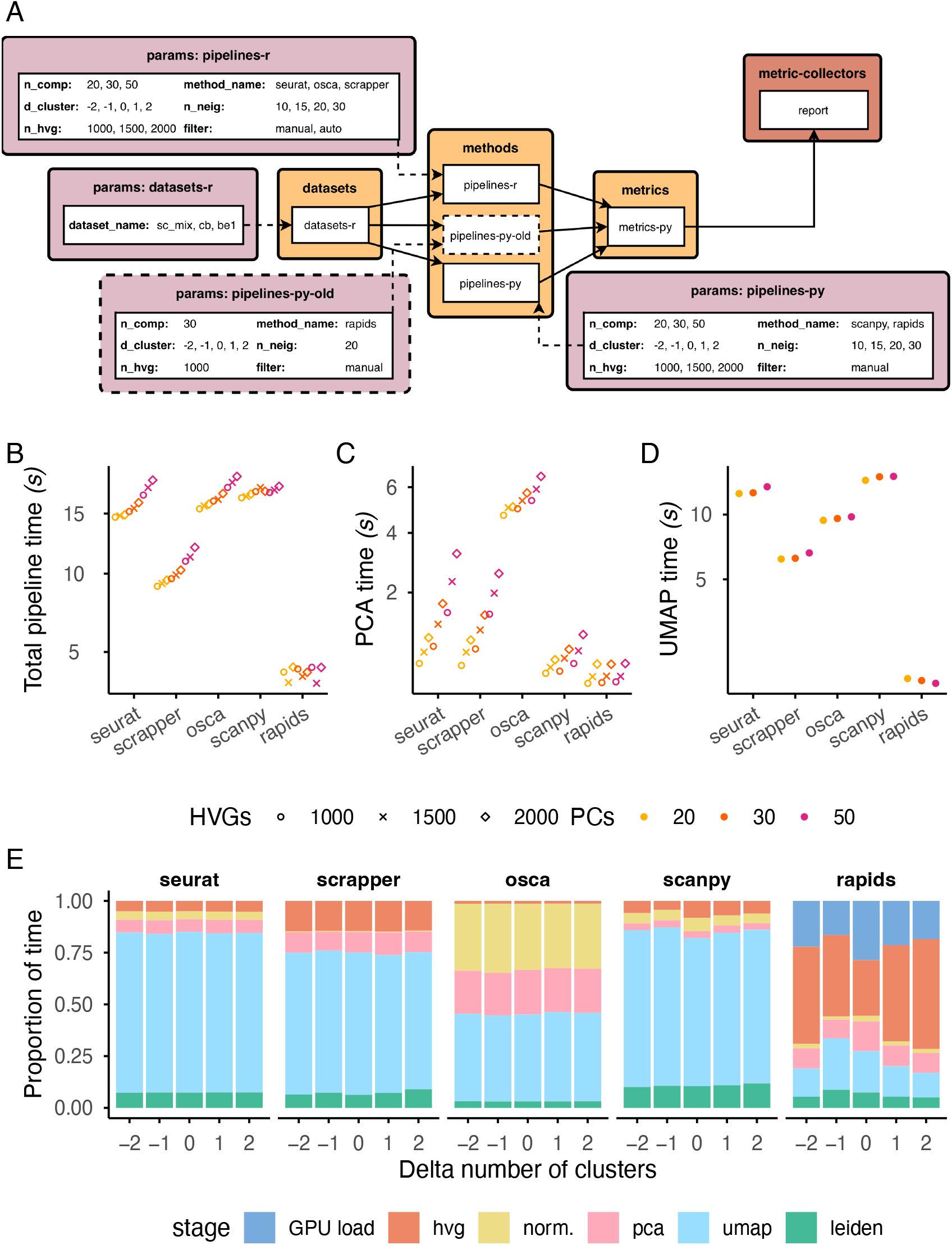
(A) Schematic of the `scrna-bench` Omnibenchmark; the full YAML file (see Methods) defines the full benchmark topology, including inputs and outputs of each stage, software environments used for each module, and pointers to repositories that contain the code that will be executed at each stage. (B), (C), and (D) give median computational times for the full pipeline (GPU load + hvg + pca + umap + leiden), PCA, and UMAP steps, respectively, for the cb dataset (7858 cells; Table S1). Note that there are multiple points for this dataset because each pipeline was run across the set of parameters shown in (A). (E) Proportion of time spent by methods in different stages of the pipeline.

While some of this amounts to a simple standardization of an existing codebase, the imposed structure also represents a vision for consistent unit testing and deeper probing of individual pipeline stages. We also built reproducible software environments in the form of (`apptainer` ) container images (see Methods). Motivated by failing to reproduce the original authors’ results using rapids (i.e., the `rapids-singlecell` package), we built two images with different versions of `rapids` . To keep the scope reasonable, we constrained the re-implementation to three datasets (`sc-mix, cb` and `be1` ) and five pipelines (`seurat, scrapper, osca, scanpy and rapids` ) from the original study. Our re-implementation adds reproducibility, standardization, and a much broader parameter set, supporting systematic benchmarking of single-cell analysis pipelines.

While orchestration via Omnibenchmark gives a basic monitoring of computational performance (memory and CPU via `snakemake` [14]), the original benchmark already captured timings for the individual steps of each pipeline. While computational cost is generally a secondary consideration, comparing them across pipelines and datasets reveals interesting trends about the implementations (Figure 1B-C and S1). For example, osca seems to have a slower PCA implementation, while `scanpy` has a slower UMAP implementation.`rapids` (a GPU implementation) and `scrapper` (a C++ library with R bindings) are the fastest of all pipelines (Figure S2). Not surprisingly, compute times are strongly associated with the parameters (e.g., number of nearest neighbors in graph building, number of features used in dimensionality reduction); see Figure 1B. The pipelines spend varying proportions of time in the multiple steps (Figure 1D). While the CPU methods were benchmarked here on single cores, many steps support multithreading. However, a more in-depth investigation of `scanpy` and `scrapper` in terms of CPU (and memory) usage along the pipeline steps highlights that `scanpy` indeed achieves noticeable performance improvements from 1 to 4 threads, though far from a 4-fold increase, and `scrapper` only achieves only modest improvements from multithreading for a dataset of this size (Figure S3). We note that runtime comparisons between CPU- and GPU-based pipelines reflect several architectural differences and should not be interpreted as hardware-independent measures of algorithmic efficiency.

The primary endpoint of the original benchmark [21] is partitions (or clusters) of cells. Although performance measures at other stages (e.g., embeddings, graphs [26]) of the pipelines were computed in `scrna-bench`, for brevity, we focus here on Adjusted Rand Index (ARI) (using the cell annotations from the original authors as ground truth) as a widely-used standard for clustering results in single-cell benchmarks [21, 27, 28], with correlation across remaining metrics reported in Figure S4.

Unsurprisingly, the number of specified clusters strongly affects the clustering performance (Figure 2A), while the dependence on the number of highly variable genes (HVGs), the number of neighbors in the nearest neighborhood graph and the number of PCA components is less clear (see Figure S5). Note that our comparisons represent complete (largely-default) pipeline configurations, so differences are not attributable to individual components. Also important to note is that there is no consistent top performer across datasets and the stability can vary greatly across different parameterizations, especially for the cb dataset (Figures S5, S8). Of further interest is whether the faster pipelines, `rapids` and `scrapper`, are able to maintain high clustering accuracy with increasing complexity in datasets; there is a notable reversal of top performance for `rapids` and `scrapper` for the `be1` and `cb` datasets (Figure 2B). In particular, suboptimal performance is exhibited by `scanpy, rapids` and `seurat` on the `be1` dataset (also discussed in [21]).

**Fig. 2.**
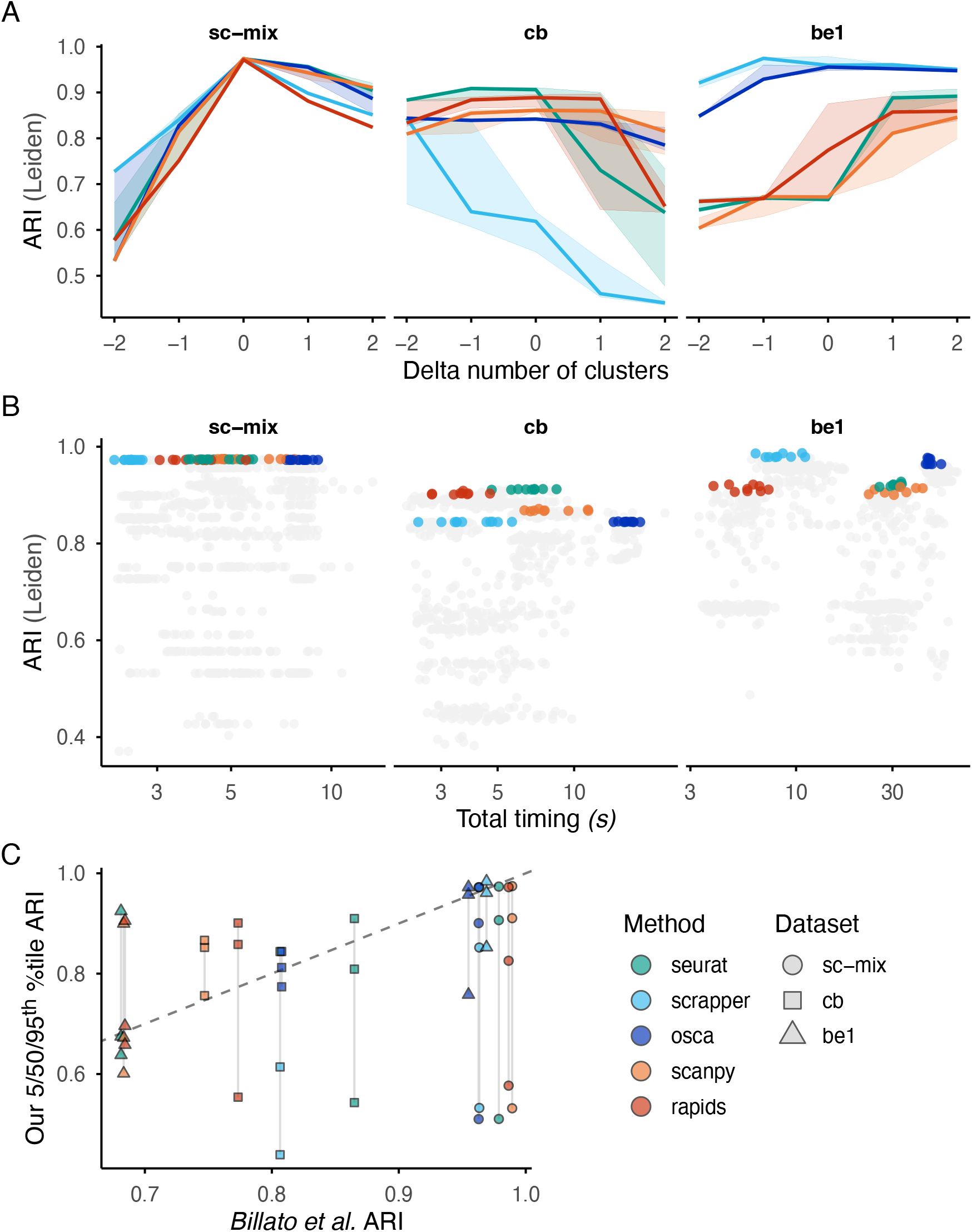
(A) Median ARIs obtained from Leiden clusters as a function of the delta number of clusters with the shaded area representing the interquartile range computed from different runs across the parameter sets: number of PCs (`n_comp` ), number of NNs (`n_neig` ), and number of selected features (`n_hvg` ). Positive and negative values of the delta number of clusters represent over- and under-clustering, respectively, relative to the expected number of clusters (cell types) in the dataset; see Methods for further description. Here, we removed the runs where the observed number of clusters did not match the specified number of clusters. (B) Tradeoff between the top 5% of Leiden ARI values (coloured) for each method within each dataset, and the corresponding runtimes (grey points represent remaining parameter combinations). (C) Comparison of the 5*/*50*/*95^*th*^ percentile ARI obtained in our analysis with those obtained in the original study [21].

Exposing a wider grid of parameters led to improvements in ARI (Figure 2C) over those reported in the original paper (see Table 1 of [21]); however, this is largely due to searching across a wider parameter space. In many cases, we can also find pipelines across the (specified) parameter space that perform worse than that reported in the original study, especially for the `cb` dataset (Figure 2C). One notable finding is that the resolution parameter is not parameterized equally across pipelines (see Figure S6B), motivating us to implement the number of clusters as the parameter of interest instead of the resolution parameter (as done in [29]).

As an alternative to visual inspection and given the now-broad parameter space investigated in `scrna-bench,` we utilized linear models to get a crude overall view of what factors affect (ARI) performance. In the first instance, we fit a standard linear model with ARI (of the Leiden clustering) as the response and the pipeline and (categorical) parameters as explanatory variables with the reference pipeline being `osca` with the correct number of clusters, 20 PCs, 1000 HVGs and 10 nearest neighbors (NNs). Thus, parameter estimates represent improvements (positive values) or worsening (negative values) of performance relative to this reference (see Figure S8); the intercept of this model represents average performance of the reference pipeline (Figure S8A; see Supp. Table S2 for the full model results). The results highlight largely dataset-specific effects.

For example, it is modestly detrimental to use a larger number of NNs for the `cb` dataset. In the `be1` dataset, some pipelines are notably worse on average than `osca` and automated filtering decreases performance slightly (also shown in Figure 2B). The model confirms that misspecifying the number of clusters stands out as a key contributor to all datasets (Figure S8 and Supp. Table S2). Fitting an analogous random effects model for each group variable (see Methods), the explained variance can be partitioned (see Figure S8B). Interestingly, most of the variation in ARI performance is still driven by the pipeline used (for datasets `cb` and `be1`) and by the number of clusters (for the “easier” `sc-mix` dataset); filtering and number of NNs explain only very small proportions of the variance and number of PCs and HVGs do not explain any variation.

Attempting to reproduce the original pipelines, we ran into an unusual case with the `sc-mix` dataset and version `0.13.5` of the `rapids` package, where the ARIs dropped significantly across all parameterizations (Figure S9). We traced this back to instabilities in the underlying PCA algorithm and implicit choice of the SVD solver (Figure S10). While we did not run into the same issue with subsequent versions, it does highlight the need for (re)testing multiple versions of an implementation, naturally leading to the perspective of benchmarks behaving like tests for packages, especially as they evolve over time. Even though the scale of this may be out of scope for package unit tests, it is feasible and offers an interesting dimension to large-scale “continuous” benchmarks where one could run the same benchmark across hardware and software environments, with different versions of the used packages.

We view `scrna-bench` as a canvas for exploring the many orthogonal directions that one could probe deeper. We evaluated “methods as pipelines”, following the canonical pipelines as much as possible. This resulted in mixing varying philosophies such as the default choice of scaling genes in `seurat, scanpy`, and `rapids`, while not doing so in `osca` and `scrapper` . A stricter evaluation of the implementations would evaluate both (and further) philosophies.

## Conclusion

We re-implemented and enhanced an arms-length benchmark by harmonizing stages, standardizing outputs and exposing parameters, while preserving most of the original code. This helps maintain reproducibility and facilitates extensibility. All artifacts from the benchmark have been archived on Zenodo and the code repositories (and therefore the entire benchmark) are reproducible, can be forked, modified, and re-run or re-designed to answer related methodological questions. We did this in part to pique interest in the community for doing such standardized studies in a more open and collaborative manner.

Our study has limitations. Currently, we only evaluate three datasets across five pipelines from the original study, and for all except `rapids`, only one software environment was tested. In particular, it was recently pointed out that the analysis implementation and version used can introduce considerable variability to downstream analyses [30]. This is not surprising since many tools are under active development and implementation tradeoffs are chosen individually by developers. An obvious extension to our analysis here is an expansion of the software environments across (recent) versions of packages (being conscious of the fact that some pipeline interfaces may change over time). Nonetheless, the flexibility required for the expansion and orchestration is already available in Omnibenchmark. CPU-based methods were tested on a single core; this seems justified given that of the 2 pipelines tested (`scanpy and scrapper` ), neither take full advantage of multi-threading.

One notable change to the design could be to expand the *stages* of the pipeline into individual steps. This has tradeoffs: it brings efficiencies because a pipeline that is identical except for the final parameter (e.g., number of clusters) was run in its entirety in `scrna-bench`, while splitting the stages into multiple steps may require storage and reading/writing of potentially large and redundant objects. However, such a reorganization would allow the scaling of the number of pipelines tested, and investigation of the interactions between the steps of a pipeline, while still allowing benchmarking “slices” that focus on specific steps. Moreover, `scrna-bench` (and that from Billato et al.: [21]) currently does not have an integration step, which is generally a requirement for multiple-sample processing. We propose to do some of this expansion as a community, because it will facilitate a consensus on the key areas of the parameter space to explore and the appropriate metrics of success. Lastly, pipelines use feature selection that focus on variability of expression and only the number of features is modulated in the current benchmark; many alternative options are available [31]. Altogether, we propose that benchmarks should become continuous entities: diversity of datasets, parameters, methods and metrics will add to the plurality of evaluation while pipelines are constantly evolving.

Overall, `scrna-bench` is a template for standardizing (single-cell) benchmarks. This strategy can be applied to a wide range of other computational tasks, guaranteeing not only reproducibility but also a well-defined benchmark topology and the corresponding artifacts (code, datasets, software environments, results) of the benchmark.

## Methods

`scrna-bench` is comprised of 4 stages: datasets, methods, metrics and a (“metric collector”) report. The code is organized into reusable *modules* that are structured within these stages. Specific behavior of these modules in response to choices, such as the number of PCA components or the number of highly variable genes (HVGs), can be injected via parameters supplied to these modules. Modules are written to be compliant with the parameterizations, and can then be orchestrated from the specifications in the benchmark plan definition. Ultimately, the benchmark plan is self-describing of all the elements that make up the benchmark (e.g., with pointers to remote repositories).

The **datasets** stage has only one module `scrna-bench/datasets-r`, written in R, which downloads the datasets and stores them as `h5ad` objects. Although modules could have been created separately per dataset, we structured the common logic into one module. The choice of dataset to be downloaded can be controlled via the `dataset_name` parameter, which is a design choice that we also followed for other modules.

Our **methods** stage has three modules, `scrna-bench/pipelines-r` (for R; `seurat, osca and scrapper` ),`scrna-bench/pipelines-py` (for Python; `scanpy` and rapids) and `scrna-bench/pipelines-py-old` (a second execution of `pipelines-py` under a different software environment; discussed below). Following a similar logic as above, we structured pipelines in one of the two modules and parameterize: the number of PCA components (`n_comp` ), number of NNs (`n_neig` ), number of HVGs (`n_hvg` ), and *δ*_cluster_, defined as the signed offset from the true number of clusters (*k*_pred_ = *k*_true_ + *δ*_cluster_; `d_cluster` ). The latter is used to iteratively search for a resolution parameter that yields the target number of clusters. In this process, the iterative search sometimes failed to yield the desired number of clusters (see Figure S7). Where necessary, we discarded such runs. QC filtering was performed either with manual, distribution-guided cutoffs or with automated cutoff selection using functions provided by some pipelines, and this choice was also parameterized. As output, this stage yields the predicted cluster labels and PCA embeddings alongside the wall-clock time for each pipeline subunit (e.g. filtering, PCA, Leiden, UMAP).

The **metrics** stage takes as input the predicted cluster labels and PCA components from the methods stage, alongside the ground-truth cell type labels from the datasets stage. Cells lacking a ground-truth annotation are excluded prior to metric computation. Metrics are computed for both Leiden and Louvain cluster assignments and fall into two categories. The first is label agreement metrics, which compare predicted cluster labels against ground-truth cell type annotations: Adjusted Rand Index (ARI), Normalized Mutual Information (NMI), Fowlkes-Mallows score, homogeneity, completeness, and V-measure. The second is cluster structure metrics, which assess the internal geometry of the clustering in PCA space without reference to ground truth: Silhouette score, Davies-Bouldin index, and Calinski-Harabasz index. In addition to these metrics, the stage records the number of clusters produced by each algorithm (which may differ from the number implied by the parameterization when the iterative search does not converge) and wall-clock timings for each pipeline subunit. The metrics computed here are finally aggregated in the metric collector (report) module.

We created three Apptainer images to encapsulate the software environments used in `scrna-bench (r_apptainer, py_apptainer_rsc_0.13.5, py_apptainer_rsc_0.14.1` ). All images are Ubuntu-based. One contains the R software stack required to run the R pipelines. The remaining two contain the Python software stack and are identical in composition except for the version of `rapids` installed. The primary benchmark is run on the image with the newer version. To assess the impact of the `rapids` update, we run a subset of the benchmark on the older image and compare results against matched runs on the newer one.

All computations were run on an Ubuntu 24.04.3 LTS server with AMD EPYC 7763 64-Core CPU Processor and NVIDIA AD104GL (L4) GPU. We did not apply any specific resource constraints.

## Linear modeling

Altogether, we fit two analogous linear models with the same response and explanatory variable, one as a standard fixed effect model and the other as a full random effects model. The model fits ARI (of the Leiden clusters) as the response, with method (pipeline) and parameters coded as categorical variables. We fit a separate model for each dataset. For the model formula in R, the standard linear model is:

~~~
ari ∼ method + d_cluster + filtering + n_comp + n_neig + n_hvg
~~~

and the random effects model (fit with the lmerTest package [32]) is:

~~~
ari ∼ (1|method) + (1|d_cluster) + (1|filtering) + (1|n_comp) + (1|n_neig) + (1|n_hvg)
~~~

We do not strictly interpret statistical significance of these estimates given that we do not have a random sample, *per se*, but the estimates and statistics give a crude estimate of which parameters are influential to (ARI) performance. Supplementary Table S1 gives a summary of the coefficients and their standard error estimates, as well as the *R*^2^ of the models (0.62-0.77).

## Resource profiling

For the in-depth resource profiling of `scanpy` and `scrapper` (Figure S3), a sub-benchmark was constructed on separate branches (`multithreading` ) of the repositories that contained 1 parameter set (dataset = `pbmc68k,` Δcluster = 0, manual filtering, 40 PCs, 15 NNs, 2000 HVGs) and 1 dataset (`pbmc68k`: [33]), and implementations that expose the maximum number of threads as a parameter. The implementations record the start and end times of each pipeline step, so they can be aligned with the `denet` [34] traces. Since each pipeline run is a single rule, `denet` ran each of the 6 `snakemake` rules (pipeline:{`scrapper,scanpy` } x threads:{1,4,8}) separately with resource usage sampled every 50 milliseconds.

## Supplementary information

We have included an excel file containing the fixed-effect linear model summaries.

## Declarations

### Ethics approval and consent to participate

Not applicable.

### Consent for publication

Not applicable.

### Availability of data and materials

Details on the datasets used in this benchmark are described in the original paper [21]. A snapshot of all benchmarking artifacts, including the code as it stood at the time of analysis (as well as software environment definitions, the `Snakefile`, all outputs of the benchmark), has been archived on Zenodo at https://doi.org/10.5281/zenodo.19886347. The primary code repositories are located on GitHub at https://github.com/scrna-bench/pipelines-plan and https://github.com/scrna-bench/pipelines-analysis. `scrna-bench/pipelines-plan` documents the benchmark plan and links to the various workflow modules, while `scrna-bench/pipelines-analysis` contains scripts for downstream analyses, figure generation, and result summaries. The software environments are defined in the planning repo and the links to the images are present in the benchmark yaml file. We also include a summary of the images, as they stand, at the time of publishing (Tables S2 and S3).

### Author contributions

AC: Conceptualization, Methodology, Software, Validation, Formal analysis, Investigation, Data curation, Methodology, Writing – Original Draft, Visualization. TK: Methodology, Software. BC: Conceptualization, Methodology, Software, Validation, Data Curation. PB: Methodology, Visualization. ME: Software, Validation. SG: Software, Validation. MK: Software, Validation, Visualization. SL: Software, Validation. IM: Conceptualization, Methodology. AM: Methodology, Software. DW: Software, Methodology. MDR: Supervision, Conceptualization, Formal analysis, Visualization. All authors: Writing – review & editing.

## Acknowledgments

We acknowledge discussions with Ilaria Billato and Davide Risso, whose original paper we are reproducing and extending, and provided the basis for our study. We also thank our hosts at Hotel Eiger in Mürren Switzerland for providing a venue for our lab retreat in which we embarked on this project.

## References

[1] Bendall, S. C. et al. Single-cell mass cytometry of differential immune and drug responses across a human hematopoietic continuum. Science 332, 687–696 (2011).

[2] Stoeckius, M. et al. Simultaneous epitope and transcriptome measurement in single cells. Nat. Methods14, 865–868 (2017).

[3] Buenrostro, J. D. et al. Single-cell chromatin accessibility reveals principles of regulatory variation. Nature523, 486–490 (2015).

[4] Dixit, A. et al. Perturb-seq: Dissecting molecular circuits with scalable single-cell RNA profiling of pooled genetic screens. Cell 167, 1853–1866.e17 (2016).

[5] Liu, C. et al. Multitask benchmarking of single-cell multimodal omics integration methods. Nat. Methods22, 2449–2460 (2025).

[6] Lim, J. et al. Advances in single-cell omics and multiomics for high-resolution molecular profiling. Exp. Mol. Med. 56, 515–526 (2024).

[7] Zappia, L., Phipson, B. & Oshlack, A. Exploring the single-cell RNA-seq analysis landscape with the scRNA-tools database. PLoS Comput. Biol. 14, e1006245 (2018).

[8] Zappia, L. & Theis, F. J. Over 1000 tools reveal trends in the single-cell RNA-seq analysis landscape.Genome Biol. 22, 301 (2021).

[9] Wolf, F. A., Angerer, P. & Theis, F. J. SCANPY: large-scale single-cell gene expression data analysis.Genome Biol. 19, 15 (2018).

[10] Amezquita, R. A. et al. Orchestrating single-cell analysis with bioconductor. Nat. Methods (2019).

[11] Lun, A. T. L. & Kancherla, J. Powering single-cell analyses in the browser with WebAssembly. J. Open Source Softw. 8, 5603 (2023).

[12] Hao, Y. et al. Dictionary learning for integrative, multimodal and scalable single-cell analysis. Nat. Biotechnol. 42, 293–304 (2024).

[13] Virshup, I., Rybakov, S., Theis, F. J., Angerer, P. & Wolf, F. A. anndata: Access and store annotated data matrices. J. Open Source Softw. 9, 4371 (2024).

[14] Mölder, F. et al. Sustainable data analysis with snakemake. F1000Res. 10, 33 (2021).

[15] Di, T. P. et al. Nextflow enables reproducible computational workflows. Nature biotechnology 35, 316–319 (2017).

[16] Luecken, M. D. et al. Defining and benchmarking open problems in single-cell analysis. Nat. Biotechnol.43, 1035–1040 (2025).

[17] Mallona, I. et al. Omnibenchmark: transparent, reproducible, extensible and standardized orchestration of solo and collaborative benchmarks. arXiv [q-bio.OT] (2026).

[18] Capella-Gutierrez, S. et al. Lessons learned: Recommendations for establishing critical periodic scientific benchmarking. bioRxiv (2017).

[19] Dicks, S. et al. GPU-accelerated single-cell analysis at scale with rapids-singlecell. arXiv [q-bio.GN] (2026).

[20] Parks, B. & Greenleaf, W. Scalable high-performance single cell data analysis with BPCells. bioRxivorg2025.03.27.645853 (2025).

[21] Billato, I. et al. Benchmarking large-scale single-cell RNA-seq analysis. bioRxivorg 2025.10.28.681564 (2025).

[22] Heumos, L. et al. Best practices for single-cell analysis across modalities. Nat. Rev. Genet. 24, 550–572 (2023).

[23] Luecken, M. D. & Theis, F. J. Current best practices in single-cell RNA-seq analysis: a tutorial. Mol. Syst. Biol. 15, e8746 (2019).

[24] Sonrel, A. et al. Meta-analysis of (single-cell method) benchmarks reveals the need for extensibility and interoperability. Genome Biol. 24, 119 (2023).

[25] Cao, Y. et al. The current landscape and emerging challenges of benchmarking single-cell methods. Brief. Bioinform. 26, bbaf380 (2025).

[26] Luo, S., Germain, P.-L., von Meyenn, F. & Robinson, M. D. On metrics for subpopulation detection in single-cell and spatial omics data. Nucleic Acids Res. 53, gkaf921 (2025).

[27] Li, Z. et al. Systematic benchmarking of computational methods to identify spatially variable genes.Genome Biol. 26, 285 (2025).

[28] Chen, X. et al. Benchmarking algorithms for spatially variable gene identification in spatial transcriptomics. Bioinformatics 41, btaf131 (2025).

[29] Sun, J. et al. Beyond benchmarking: an expert-guided consensus approach to spatially aware clustering.Nat. Methods 23, 1814–1826 (2026).

[30] Rich, J. M. et al. The impact of package selection and versioning on single-cell RNA-seq analysis. Cell Syst. 101560 (2026).

[31] Pullin, J. M. & McCarthy, D. J. A comparison of marker gene selection methods for single-cell RNA sequencing data. Genome Biol. 25, 56 (2024).

[32] Kuznetsova, A., Brockhoff, P. B. & Christensen, R. H. B. LmerTest package: Tests in linear mixed effects models. J. Stat. Softw. 82, 1–26 (2017).

[33] Zheng, G. X. Y. et al. Massively parallel digital transcriptional profiling of single cells. Nat. Commun. 8, 14049 (2017).

[34] Carrillo, B. & Mallona, I. denet, a lightweight command-line tool for process monitoring in benchmarking and beyond. arXiv [cs.PF] (2025).

[35] Tian, L. et al. Benchmarking single cell RNA-sequencing analysis pipelines using mixture control experiments. Nat. Methods 16, 479–487 (2019).

[36] Arigoni, M. et al. A single cell RNAseq benchmark experiment embedding “controlled” cancer heterogeneity. Sci. Data 11, 159 (2024).

