## Supplementary Information for "Building computational benchmarks: an Omnibenchmark reimplementation of a single-cell preprocessing pipeline evaluation"

#### Supplementary Tables

| Dataset | Description | Cells | Genes | Classes | Annotation Source | Technology |
|---|---|---|---|---|---|---|
| sc-mix [35] | Five human lung adenocarcinoma cell lines, mixed in equal proportions | 3,918 | 11,786 | 5 | computational de-multiplexing | 10x Chromium |
| cb [2] | Cord blood sample with surface protein markers for class annotation | 7,858 | 20,400 | 8 | gating of ADT surface protein markers | CITE-seq |
| bel [36] | Lung cancer cell lines expressing different driver genes | 29,606 | 36,753 | 8 | cell lines tagged before multiplexing | CellPlex (10x Genomics) |
| pbmc68k [33] | Peripheral blood mononuclear cells | 68,579 | 32,738 | 8 | correlation with purified cell populations | GemCode (10x Genomics) |

**Table S1 Datasets used in this work.** Note that pbmc68k was only used for the multithreading analysis.

**Table S2** Python environments: images `py-rsc-0.13.5-latest-amd64`, `py-rsc-0.14.1-latest-amd64` from https://quay.io/repository/omnibenchmark/scrna-bench?tab=tags. The two image definitions are identical upto a difference in the version of rapids-singlecell. The shared values span both columns.

| | `py-rsc-0.13.5` | `py-rsc-0.14.1` |
|---|---|---|
| *Image* | | |
| Definition | `py.def` | |
| SIF SHA-256 | `6d7c94f32017` | `2b55605477f9` |
| *System* | | |
| OS | Ubuntu 24.04.4 LTS | |
| glibc | 2.39 | |
| Python | 3.12.12 | |
| CUDA runtime | 12.9.79 | |
| cuBLAS | 12.9.1.4 | |
| NCCL | 2.29.3 | |
| BLAS (numpy) | OpenBLAS 0.3.27 (numpy wheel) | |
| *Packages* | | |
| rapids-singlecell | **0.13.5** | **0.14.1** |
| scanpy | 1.10.4 | |
| anndata | 0.11.3 | |
| numpy | 2.1.3 | |
| scipy | 1.15.2 | |
| pandas | 2.2.3 | |
| scikit-learn | 1.6.1 | |
| numba | 0.61.0 | |
| llvmlite | 0.44.0 | |
| h5py | 3.13.0 | |
| python-igraph | 0.11.8 | |
| leidenalg | 0.10.2 | |
| louvain | 0.8.2 | |
| umap-learn | 0.5.7 | |
| pynndescent | 0.5.13 | |
| statsmodels | 0.14.4 | |
| networkx | 3.4.2 | |
| matplotlib | 3.10.1 | |
| seaborn | 0.13.2 | |
| cupy | 13.6.0 | |
| cudf | 26.2.1 | |
| cuml | 26.2.0 | |
| cugraph | 26.2.0 | |
| rmm | 26.2.0 | |
| dask | 2026.1.1 | |

**Table S3** R environment: image `r-bioc-320-latest-amd64` from https://quay.io/repository/omnibenchmark/scrna-bench?tab=tags.

| | | | |
|---|---|---|---|
| *Image* | | | |
| Definition | `r-bioc-320.def` | | |
| SIF SHA-256 | `cde733cf1e75` | | |
| *System* | | | |
| OS | Ubuntu 24.04.1 LTS | | |
| glibc | 2.39 | | |
| R | 4.4.2 | | |
| BLAS | OpenBLAS (pthread) | | |
| LAPACK | OpenBLAS 0.3.26 (pthread) | | |
| *Packages* | | | |
| BiocVersion | 3.20.0 | rARPACK | 0.11-0 |
| RSpectra | 0.16-2 | SingleCellExperiment | 1.28.1 |
| zellkonverter | 1.16.0 | BiocSingular | 1.22.0 |
| here | 1.0.2 | HDF5Array | 1.34.0 |
| mbkmeans | 1.22.0 | ggplot2 | 4.0.1 |
| scater | 1.34.1 | scran | 1.34.0 |
| BiocParallel | 1.40.2 | DelayedMatrixStats | 1.28.1 |
| rhdf5 | 2.50.2 | MultiAssayExperiment | 1.32.0 |
| SingleCellMultiModal | 1.18.0 | scuttle | 1.16.0 |
| AnnotationDbi | 1.68.0 | mclust | 6.1.2 |
| bluster | 1.16.0 | leidenbase | 0.1.36 |
| DelayedArray | 0.32.0 | dplyr | 1.1.4 |
| Seurat | 5.4.0 | patchwork | 1.3.2 |
| scrapper | 1.0.3 | TENxBrainData | 1.26.0 |
| EnsDb.Hsapiens.v75 | 2.99.0 | biomaRt | 2.62.1 |
| DropletUtils | 1.26.0 | argparse | 2.3.1 |
| GEOquery | 2.74.0 | Azimuth | 0.5.0 |
| BPCells | 0.3.1 | stringi | 1.8.7 |
| SeuratObject | 5.3.0 | SeuratDisk | 0.0.0.9021 |

### Supplementary Figures

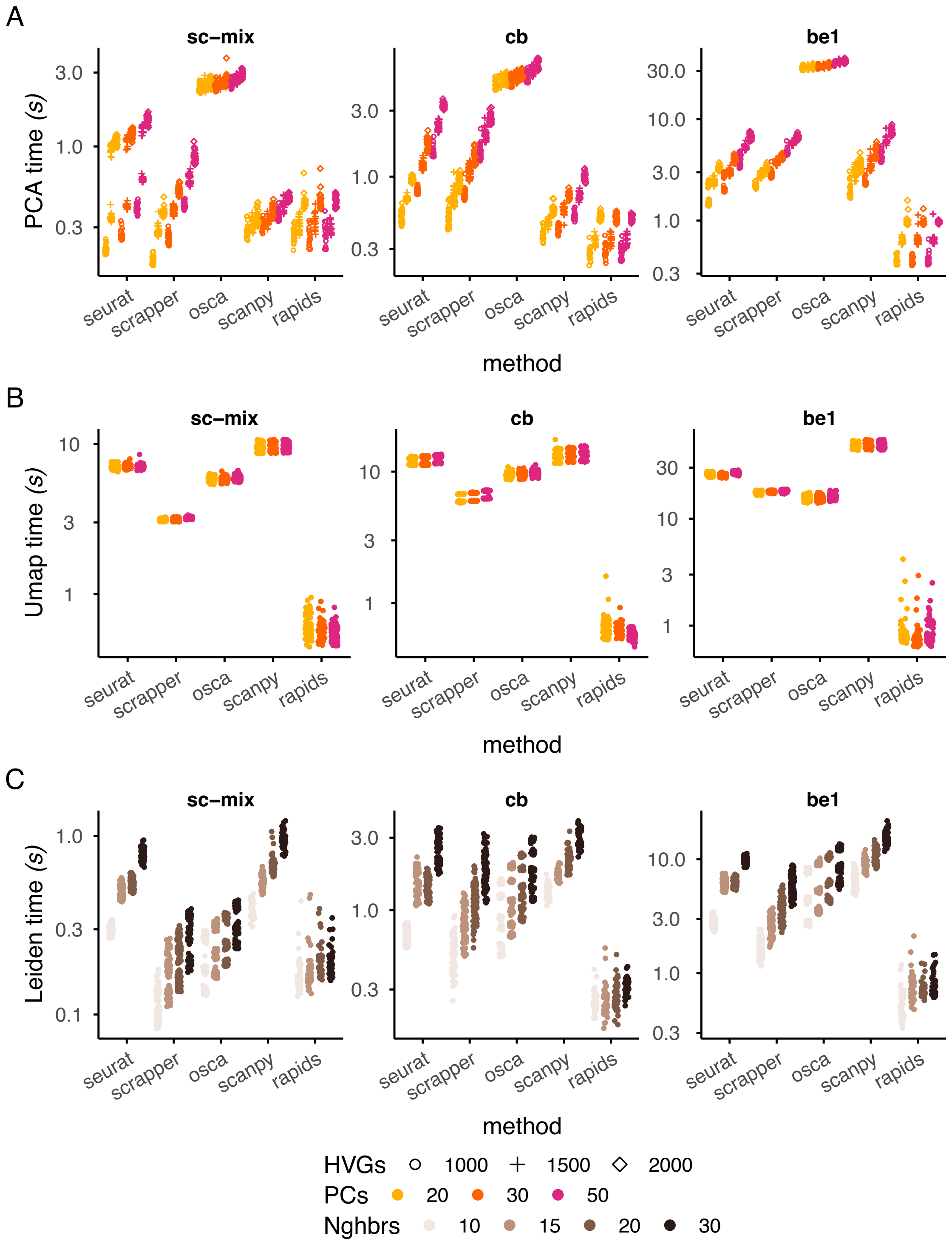


**Fig. S1 Runtime breakdown for computationally intensive pipeline stages.** Runtime (seconds) for (A) PCA, (B) UMAP and (C) Leiden clustering across the different methods and datasets. Each point represents one parameter combination; colour encodes the number of PCs (A, B) or neighbors (C), and point shape encodes the number of HVGs (A).

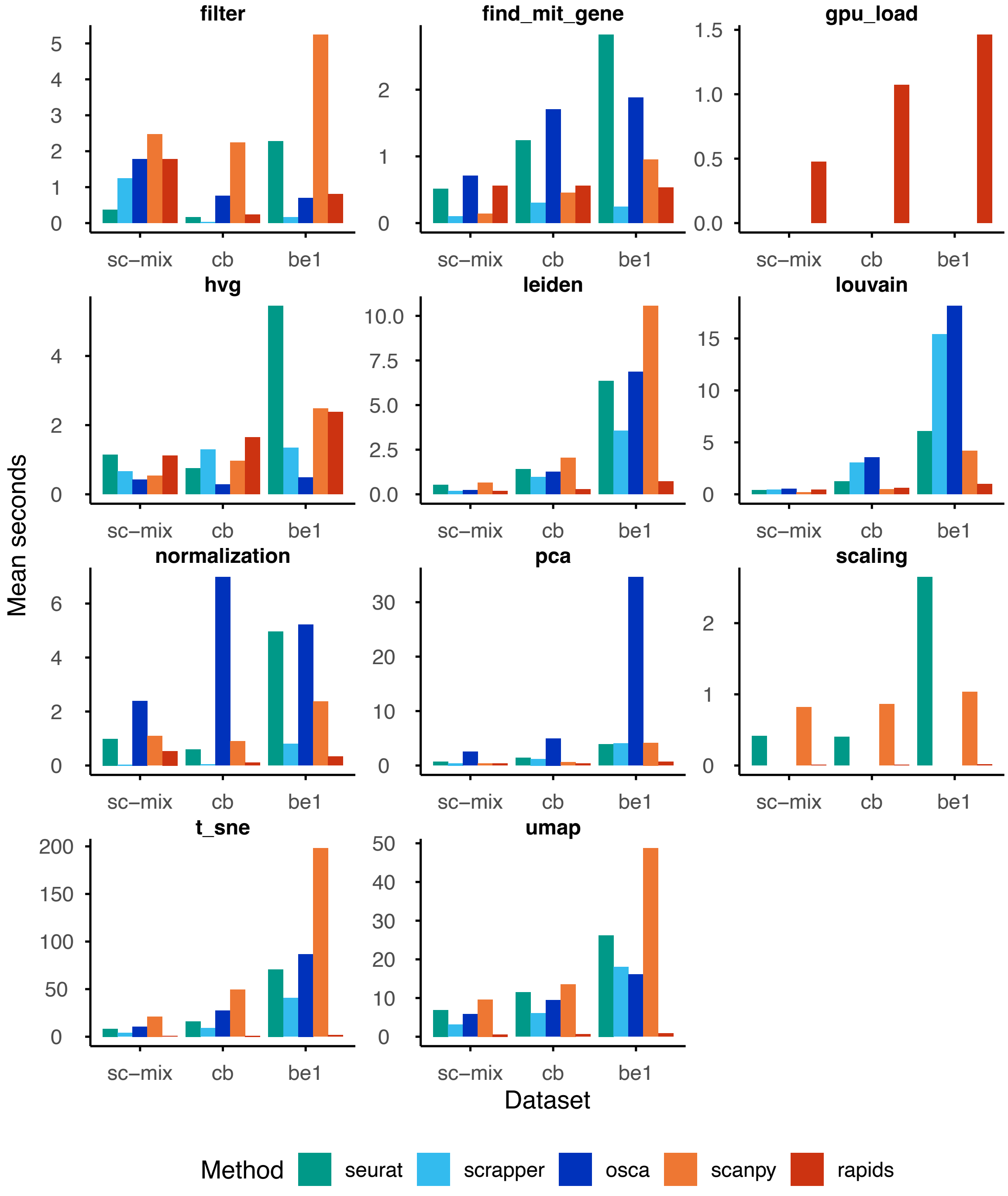


**Fig. S2 Mean runtime per pipeline stage across methods and datasets.** Bar charts show the mean execution time (seconds) for each step of the scRNA-seq analysis pipeline – covering quality control (filter, find_mit_gene), preprocessing (normalization, scaling, hvg), dimensionality reduction (pca), graph-based clustering (leiden, louvain), and visualization (t_sne, umap). The gpu_load panel reflects the initialization overhead specific to `rapids`.

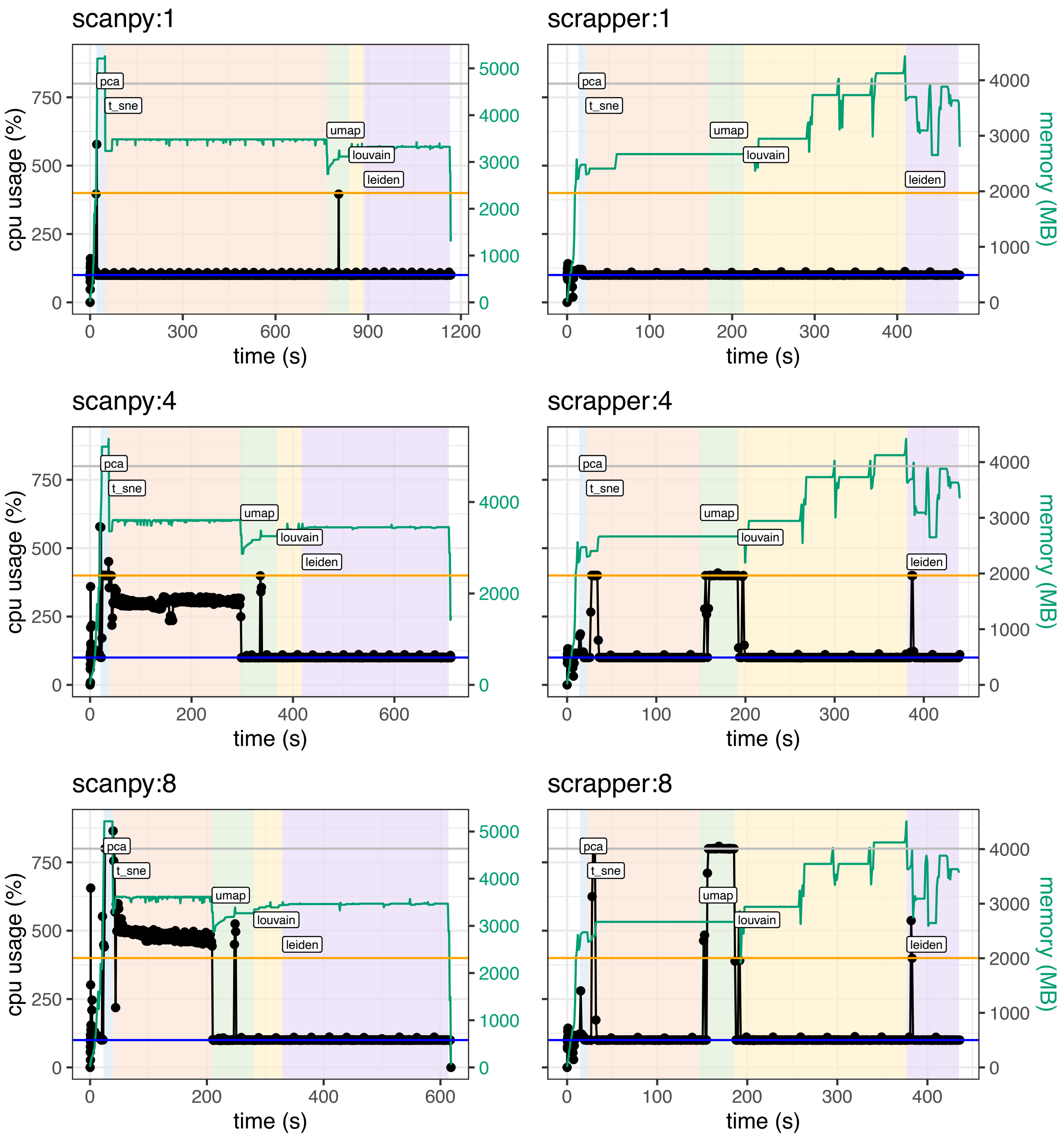


**Fig. S3 Resource usage of `scanpy` and `scrapper` when allowing multithreading.** Shown are CPU (black lines/dots) and memory usage (green lines) across the time frame of a pipeline (`denet` was configured to sample every 50 milliseconds). For both pipelines, the same parameter set was used: dataset = `pbmc68k`, $\Delta$cluster = 0, manual filtering, 40 PCs, 15 NNs, 2000 HVGs; in addition, the maximum number of threads was specified (1: top panels, 4: middle panels, 8: bottom panels). Horizontal lines are added at 100% (blue), 400% (orange), and 800% (grey) CPU usage. Coloured regions regions represent the time window of a pipeline step; labels (pca, t_sne, umap, louvain, leiden) are added to the left of each region. Note that louvain and leiden regions are comprised of multiple iterations to find a resolution that yields the correct number of clusters (all `scanpy` runs require 5 iterations, while all `scrapper` runs require 4 iterations for these parameter settings).

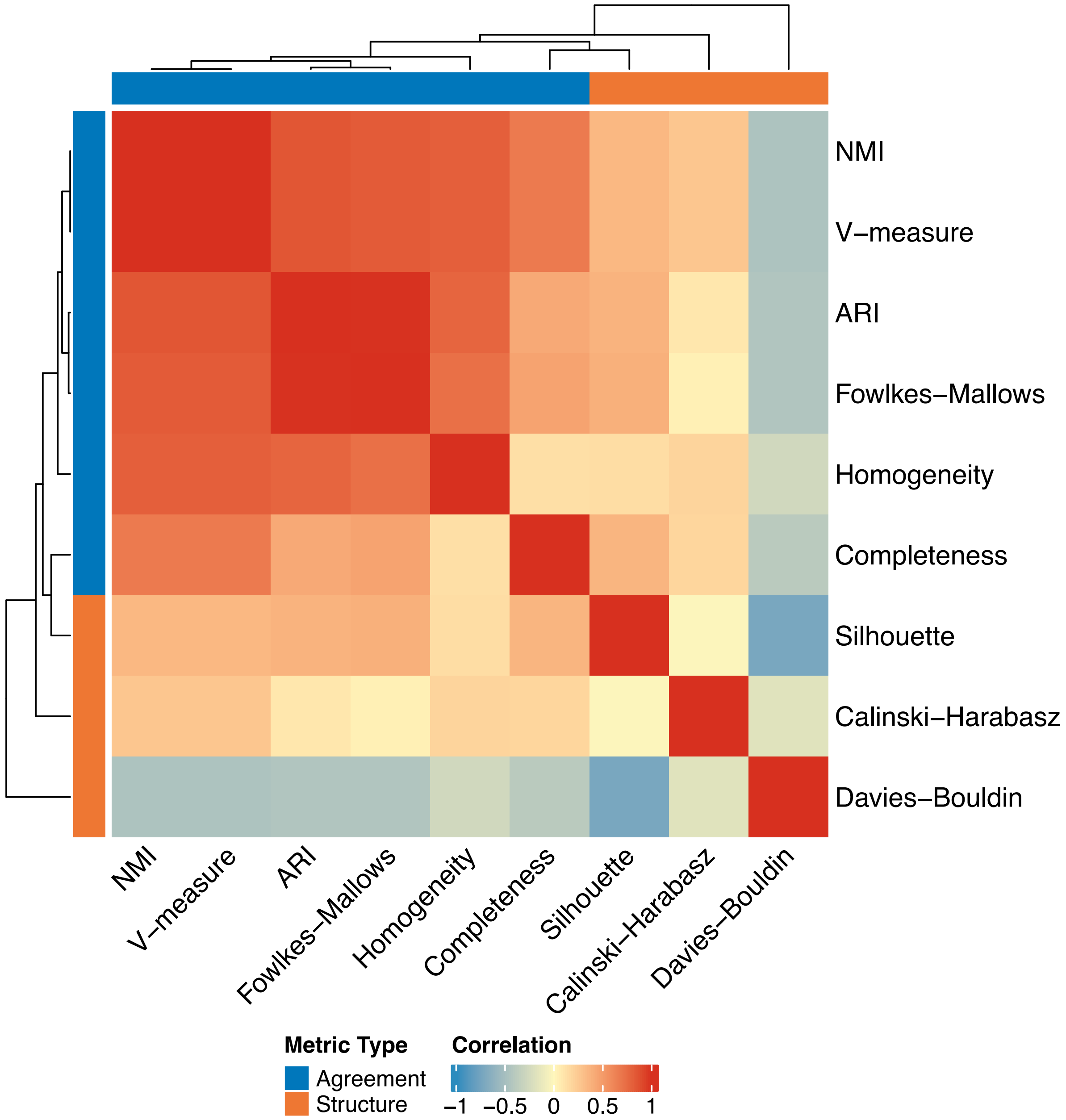


**Fig. S4 Pairwise correlations among clustering quality metrics** Spearman correlation matrix of nine clustering quality metrics computed across all pipeline runs using Leiden clustering. Metrics are grouped into agreement-based measures (ARI, NMI, Fowlkes-Mallows, Homogeneity, Completeness, V-measure), which compare predicted clusters to reference labels, and structure-based measures (Silhouette, Davies-Bouldin, Calinski-Harabasz), which assess cluster geometry without reference labels.

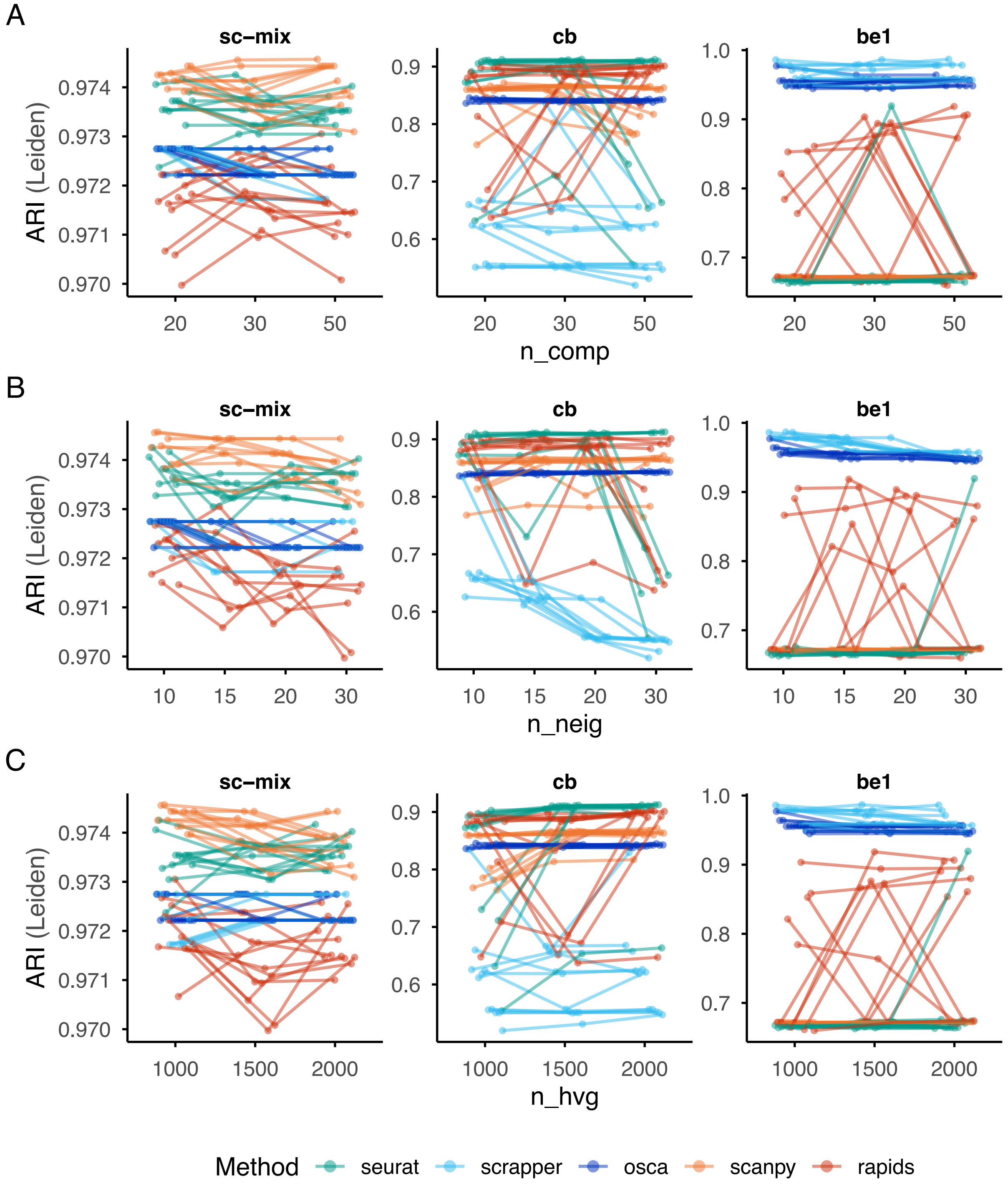


**Fig. S5 Effect of parameter choices on clustering agreement** Leiden-based ARI across varying (A) number of PCA components (`n_comp`: 20, 30, 50), (B) number of neighbors (`n_neig`: 10, 15, 20, 30), and (C) number of highly variable genes (`n_hvg`: 1000, 1500, 2000), shown for five methods across three datasets. Here, we removed the runs where the observed number of clusters did not match the specified number of clusters. Each line traces the ARI of a single parameter combination as the focal parameter varies, with all other parameters held fixed (manually filtered cells, no cluster offset).

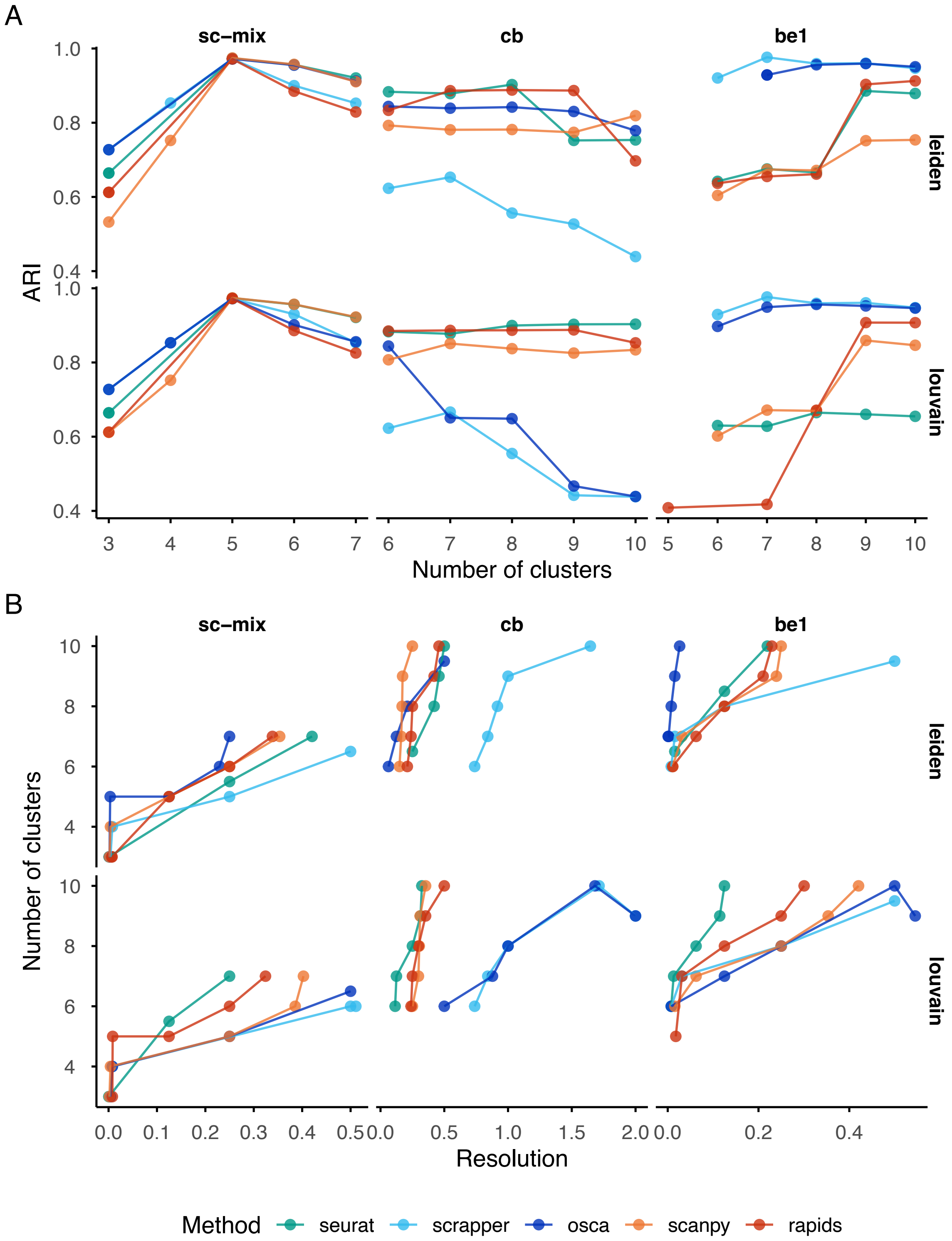


**Fig. S6 Relationships between ARI, number of clusters and resolution** Here, we examine ARI while fixing the number of PCA components (`n_comp`) to 50, number of neighbors (`n_neig`) to 20 and the number of highly variable genes (`n_hvg`) to 1000. (A) tracks the relationship between ARI and the number of clusters for the different methods. The observed differences between methods are in line with what [21] report. (B) shows the relationship between the number of clusters and the resolution.

.

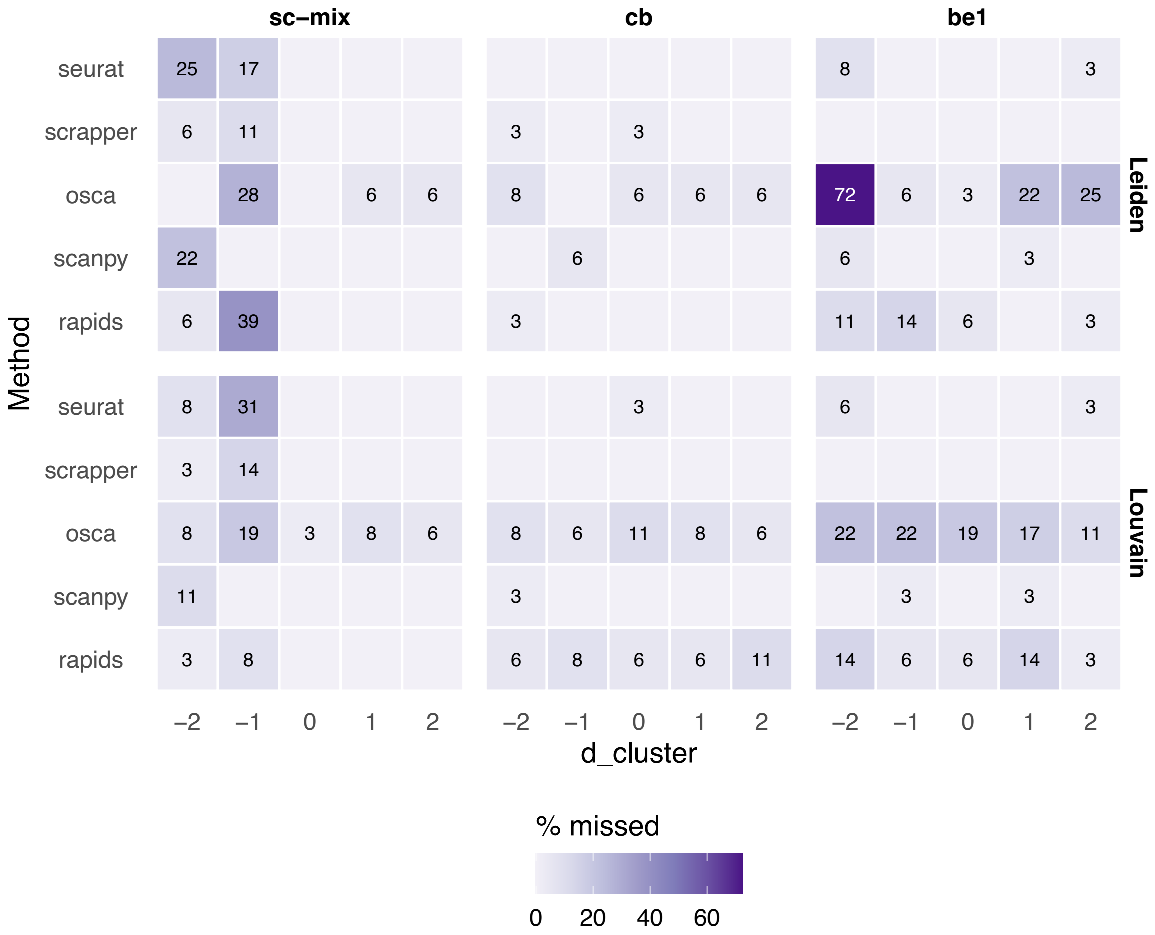


**Fig. S7 Iterative clustering misses** Plotted are how often achieved number of clusters differed from the specified number of clusters. Each square represents the percentage of missed runs from 36 runs obtained via crossing 3 values for the number of PCA components (`n_comp`: 20, 30, 50), 4 values for number of neighbors (`n_neig`: 10, 15, 20, 30), and 3 values for the number of highly variable genes (`n_hvg`: 1000, 1500, 2000)

.

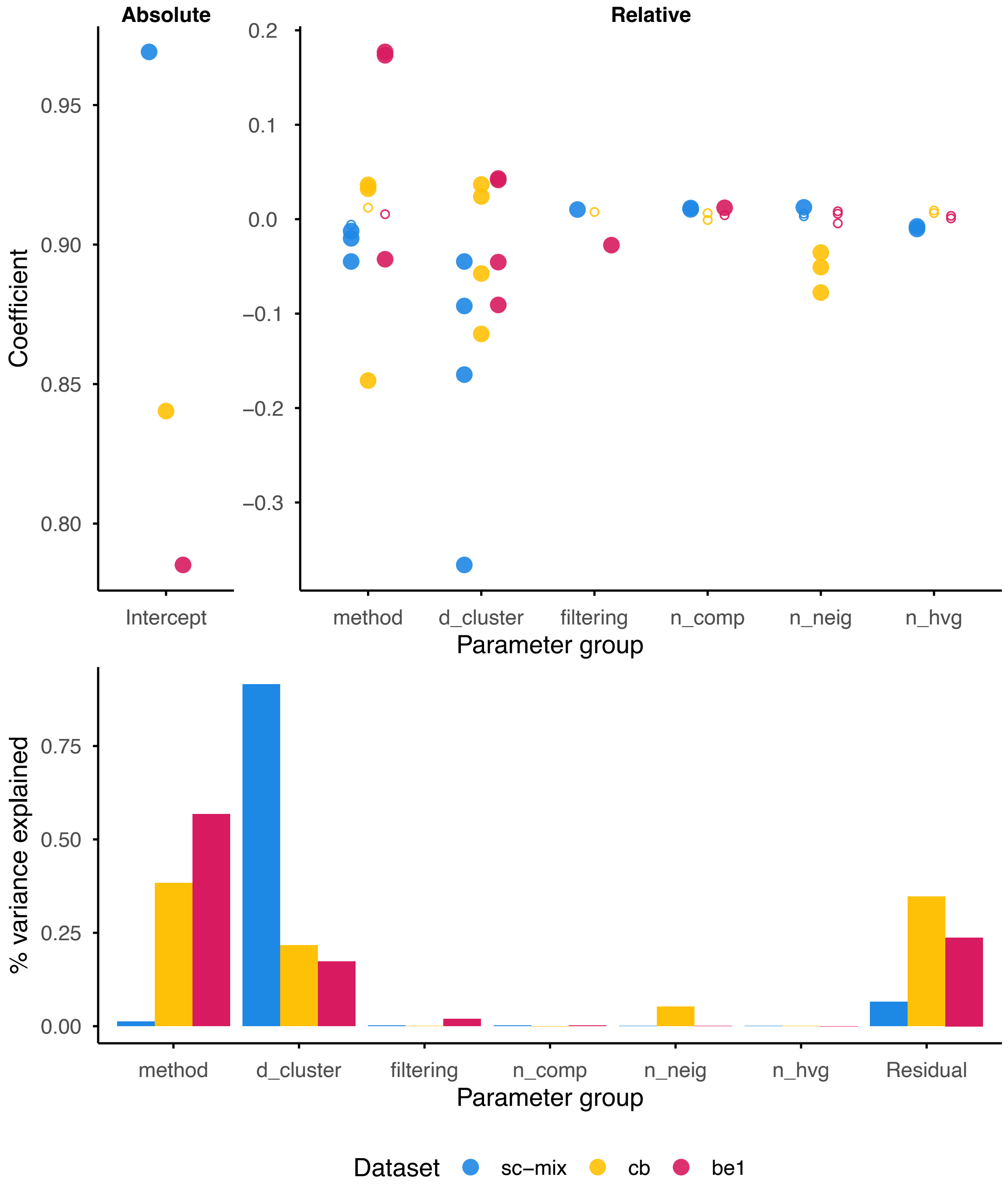


**Fig. S8 Using linear or random effects models to understand benchmark results**. (A) Plotted are the parameter estimates for a standard linear model for each dataset (linear models are fit separately for each dataset; see Methods). Interpretation of parameters is split: the "Intercept" parameter represents the average ARI under the arbitrarily-selected reference pipeline (`osca`, 20 PCs, 1000 HVGs and 10 NNs) whereas all the other parameters represent relative improvements (or deterioration) in performance. (B) Percent variance explained in an analogous random effects model (see Methods). The analysis was done after removing the runs where the observed number of clusters did not match the specified number of clusters.

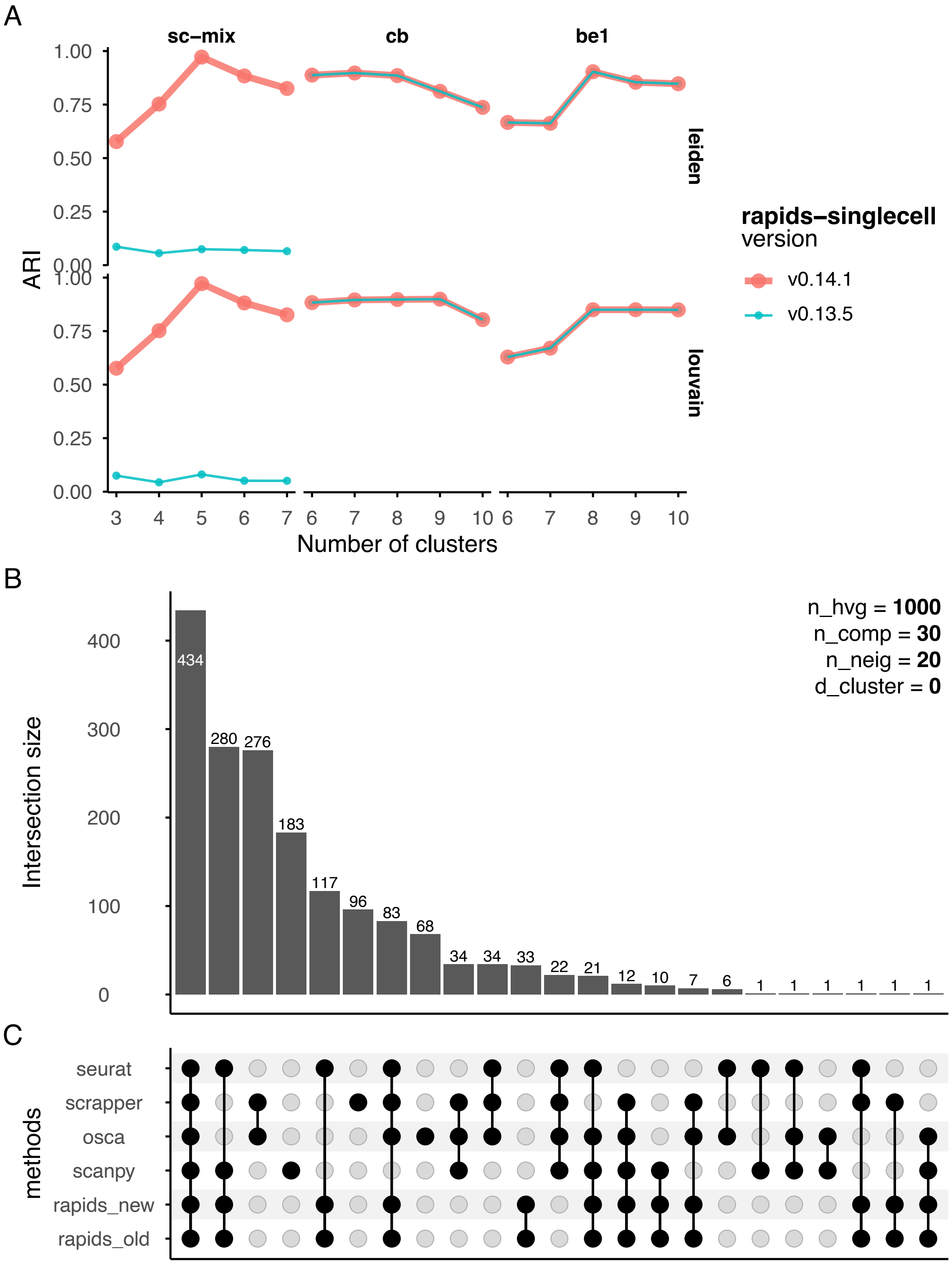


**Fig. S9 Investigation of `rapids`** For a fixed setting of the number of components (`n_comp`), number of neighbors (`n_neig`) and number of highly variable genes (`n_hvg`), we study two versions of the `rapids` package. In (A), we observe that while they mostly agree on other datasets, there is a stark drop in ARIs for version 0.13.5 in `sc-mix`. We hypothesize that this is related to the choice of the SVD solver of the PCA algorithm. To isolate this, we study an upset plot of HVGs across methods in (B). In (C), we observe that both versions are identical on the set of HVGs used for downstream analysis. This points us towards inconsistencies in the underlying PCA method, which we investigate in Figure S10.

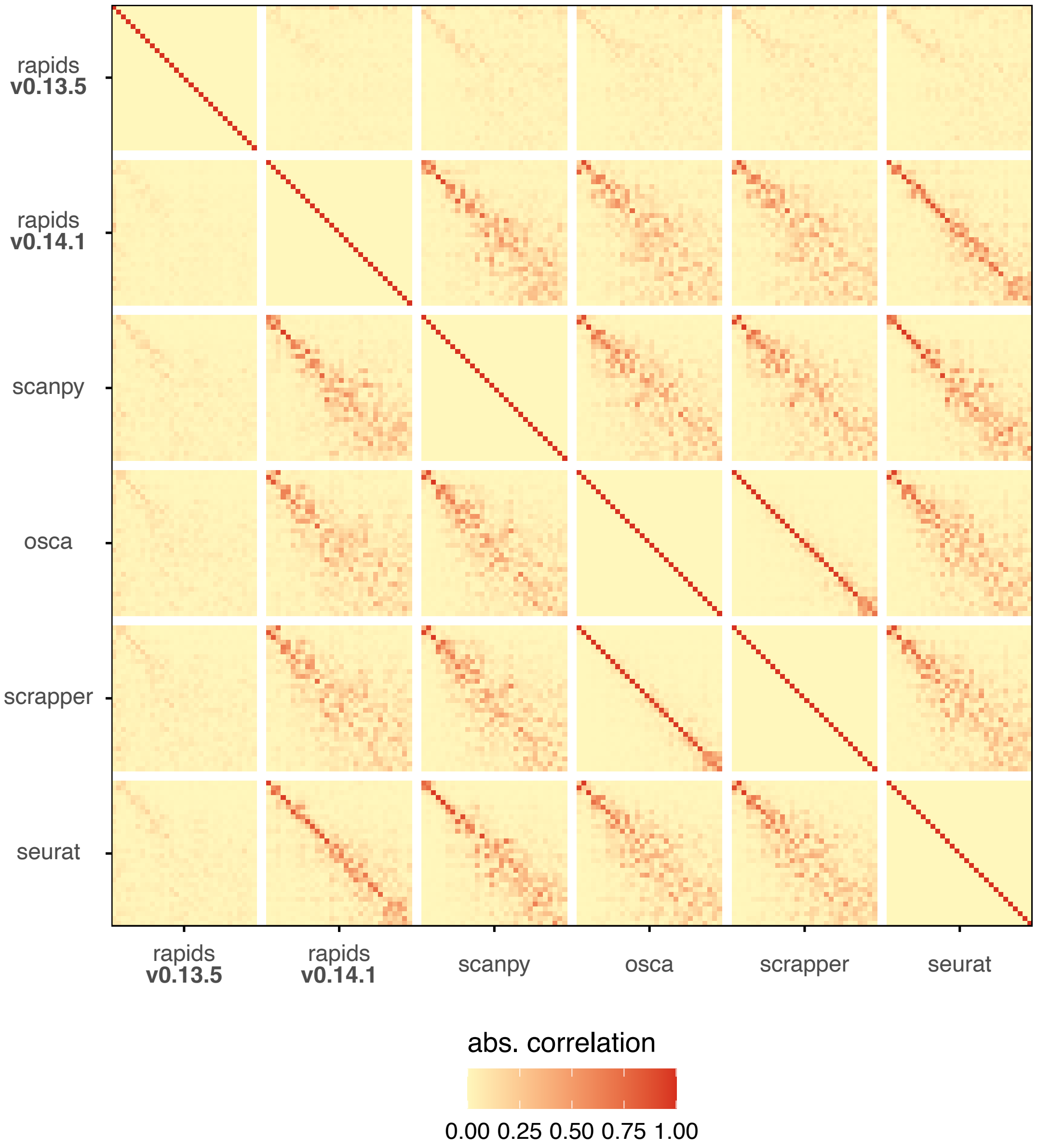


**Fig. S10 PCA correlations across methods** On the same fixed settings as S9, we look at the absolute values of the PCA correlation matrix across different methods for the `sc-mix` dataset.
